# Orthogonal outlier detection and dimension estimation for improved MDS embedding of biological datasets

**DOI:** 10.1101/2023.02.13.528380

**Authors:** Wanxin Li, Jules Mirone, Ashok Prasad, Nina Miolane, Carine Legrand, Khanh Dao Duc

## Abstract

Conventional dimensionality reduction methods like Multidimensional Scaling (MDS) are sensitive to the presence of orthogonal outliers, leading to significant defects in the embedding. We introduce a robust MDS method, called DeCOr-MDS (Detection and Correction of Orthogonal outliers using MDS), based on the geometry and statistics of simplices formed by data points, that allows to detect orthogonal outliers and subsequently reduce dimensionality. We validate our methods using synthetic datasets, and further show how it can be applied to a variety of large real biological datasets, including cancer image cell data and human microbiome project data.

## 1 Introduction

Multidimensional scaling (MDS) is a commonly used and fast method of data exploration and dimension reduction, with the unique capacity to take non-euclidean dissimilarities as its input. However, sensitivity to outliers is a major drawback [1, 2]. As arbitrary removal of outliers is undesirable, a possible alternative is to detect outliers and accommodate their influence on the MDS embedding, thus leveraging the information contained in outlying points.

Outlier detection has been widely used in biological data. Sheih and Yeung proposed a method using principal component analysis (PCA) and robust estimation of Mahalanobis distances to detect outlier samples in microarray data [3]. Chen *et al*. reported the use of two PCA methods to uncover outlier samples in multiple simulated and real RNA-seq data [4]. Outlier influence can be mitigated depending on the specific type of outlier. In-plane outliers and bad leverage points can be harnessed using ℓ_1_-norm [5–7], correntropy or M-estimators [8]. Outliers which violate the triangular inequality can be detected and corrected based on their pairwise distances [2]. Orthogonal outliers are another particular case, where outliers have an important component, orthogonal to the hyperspace where most data is located.

Although MDS is known to be very sensitive to such orthogonal outliers [9, 10], none of the existing methods addresses orthogonal outliers to the best of our knowledge. We present here a robust MDS method, called *DeCOr-MDS*, **De**tection and **C**orrection of **Or**thogonal outliers using **MDS**. *DeCOr-MDS* takes advantage of geometrical characteristics of the data to reduce the influence of orthogonal outliers, and estimate the dimension of the dataset. Our paper is organized as follows. We first describe the procedure and its implementation in detail. We then validate our method on synthetic data to confirm the accuracy and characterize the importance of different parts of our procedure. We further run the method on different experimental datasets, from single cell images and microbiome sequencing, illustrating how it can be broadly applied to interpret and improve the performance of MDS on biological datasets. Finally, we discuss the advantages and limitations of our method and future directions.

## 2 Material and Methods

### 2.1 Background: Height and Volume of n-simplices

We recall some geometric properties of simplices, which our method is based on. For a set of *n* points (*x*_1_, …, *x_n_*), the associated *n*-simplex is the polytope of vertices (*x*_1_, …, *x_n_*) (a 3-simplex is a triangle, a 4-simplex is a tetrahedron and so on). The height *h*(*V_n_, x*) of a point *x* belonging to a *n*-simplex *V_n_* can be obtained as [11]

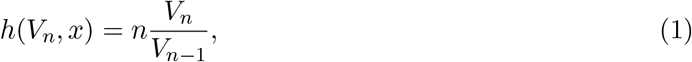

where *V_n_* is the volume of the *n*-simplex, and *V*_*n*–1_ is the volume of the (*n* – 1)-simplex obtained by removing the point *x*. *V_n_* and *V*_*n*–1_ can be computed using the pairwise distances only, with the Cayley-Menger formula [11]:

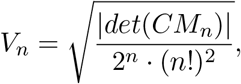

where *det*(*CM_n_*) is the determinant of the Cayley-Menger matrix *CM_n_*, that contains the pairwise distances *d_i,j_* = ||*x_i_* – *x_j_*||, as

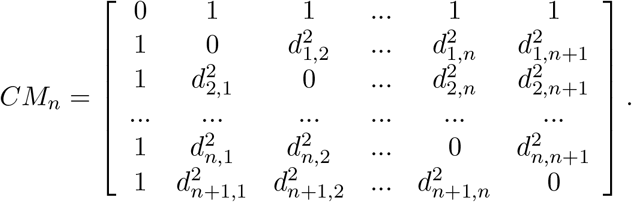

### 2.2 Orthogonal outlier detection and dimensionality estimation

We now consider a dataset **X** of size *N* × *d*, where *N* is the sample size and d the dimension of the data. We associate with **X** a matrix **D** of size *N* × *N*, which represents all the pairwise distances between observations of **X**. We also assume that the data points can be mapped into a vector space with *regular observations* that form a *main* subspace of unknown dimension d* with some small noise, and additional *orthogonal outliers* of relatively large orthogonal distance to the main subspace (Fig.1A). Our proposed method aims to infer from **D** the dimension of the main data subspace *d*\*, using the geometric properties of simplices with respect to their number of vertices: Consider a (*n*+2)-simplex containing a data point *x_i_* and its associated height, that can be computed using equation (1) in section 2.1. When *n < d*\* and for *S* large enough, the distribution of heights obtained from different simplices containing *x_i_* remains similar, whether *x_i_* is an orthogonal outlier or a regular observation (see Fig.1B). In contrast, when *n* ≥ *d*\*, the median of these heights approximately yields the distance of *x_i_* to the main subspace (Fig.1C). This distance should be significantly larger when *x_i_* is an orthogonal outlier, compared with regular points, for which these distances are tantamount to the noise.

**Figure 1:**
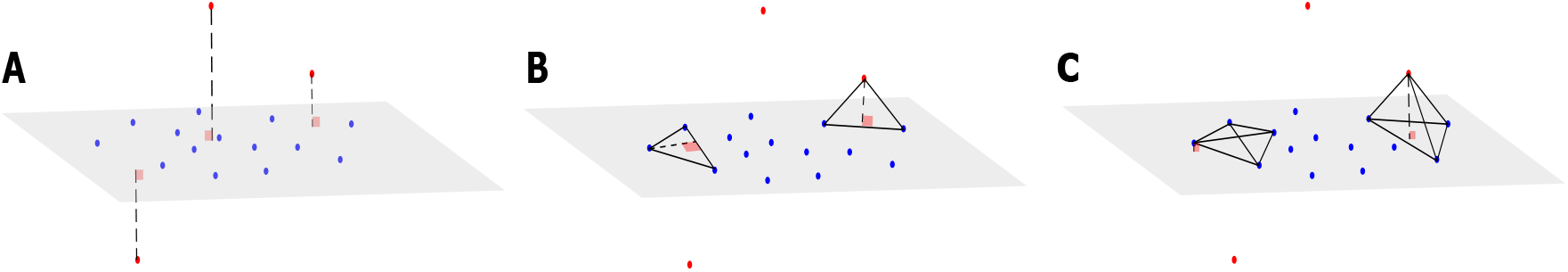
Example of a dataset with orthogonal outliers and n-simplices. **A:** We represent a dataset with regular data points (blue) belonging to a main subspace of dimension 2 with some noise, and orthogonal outliers in the third dimension. **B:** We show two 3-simplices (triangles), one with only regular points (left) and the other one containing one outlier (right). The height drawn from the outlier is similar to the height of the regular triangle. **C:** Upon adding other regular points to get tetrahedrons (4-simplices), the height drawn from the outlier gets significantly larger than the height drawn from the same point as in **(B)**.

To estimate *d*\* and for a given dimension *n* tested, we thus randomly sample, for every *x_i_* in **X**, *S*(*n* + 2)-simplices containing *x_i_*, and compute the median of the heights 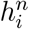 associated with these S simplices. Upon considering, as a function of the dimension *n* tested, the distribution of median heights 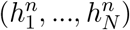 (with 1 ≤ *i* ≤ *N*), we then identify *d*\* as the dimension at which this function presents a sharp transition towards a highly peaked distribution at zero. To do so, we compute 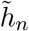, as the mean of 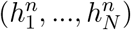, and estimate *d*\* as

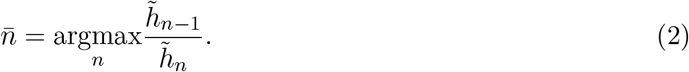

Furthermore, we detect orthogonal outliers using the distribution obtained in 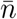, as the points for which 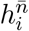 largely stands out from 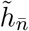. To do so, we compute 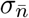 the standard deviation observed for the distribution 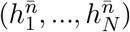, and obtain the set of orthogonal outliers **O** as

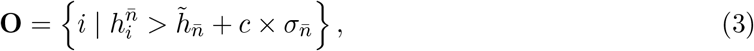

where *c* > 0 is a parameter set to achieve a reasonable trade-off between outlier detection and false detection of noisy observations.

### 2.3 Correcting the dimensionality estimation for a large outlier fraction

The method presented in the previous section assumes that at dimension *d*\*, the median height calculated for each point reflects the distance to the main subspace. This assumption is valid when the fraction of orthogonal outliers is small enough, so that the sampled *n*-simplex likely contains regular observations only, aside from the evaluated point. However, if the number of outliers gets large enough so that a significant fraction of *n*-simplices also contains outliers, then the calculated heights would yield the distance between *x_i_* and an outlier-containing hyperplane, whose dimension is larger than a hyperplane containing only regular observations. The apparent dimensionality of the main subspace would thus increase and generates a positive bias on the estimate of *d*\*.

Specifically, if **X** contains a fraction of *p* outliers, and if we consider *o_n,p,N_* the number of outliers drawn after uniformly sampling *n* + 1 points (to test the dimension *n*), then *o_n,p,N_* follows a hypergeometric law, with parameters *n* + 1, the fraction of outliers *p* = *N_o_/N*, and *N*. Thus, the expected number of outliers drawn from a sampled simplex is (*n* + 1) × *p*. After estimating 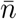 (from section 2.2), and finding a proportion of outliers 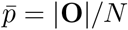 using equation (3), we hence correct 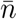 *a posteriori* by substracting the estimated bias *δ*, as the integer part of the expectation of *o_n,p,N_*, so the debiased dimensionality estimate *n*\* is

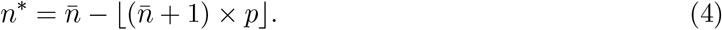

### 2.4 Outlier distance correction

Seeing that outliers have been identified previously, and because the detected dimension *d*\* is a key information on the main subspace containing regular points, it is meaningful to correct the pairwise distances that contain outliers in the matrix **D**, in order to apply a MDS that projects the outliers in the main subspace. In the case where the original coordinates cannot be used (e.g, as a result of some transformation or if the distance is non Euclidean), we perform the two following steps: (*i*) We first apply a MDS on **D** to place the points in a euclidean space of dimension *d*, as a new matrix of coordinates 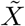. (*ii*) We run a PCA on the full coordinates of the estimated set of regular data points (i.e. 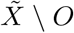), and project the outliers along the first 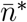 principal components of the PCA, since these components are sufficient to generate the main subspace. Using the projected outliers, we accordingly update the pairwise distances in **D** to obtain the corrected distance matrix **D**\*. Note that in the case where **D** derives from a euclidean distance between the original coordinates, we can skip step (*i*), and directly run step (*ii*) on the full coordinates of the estimated set of regular data points.

### 2.5 Overall procedure and implementation

The overall procedure, called DeCOr-MDS, is described in Algorithm 1. We also provide an implementation in Python 3.8.10 available on this [github repository](https://github.com/wxli0/DeCOr-MDS).

#### Algorithm 1 DeCOr-MDS

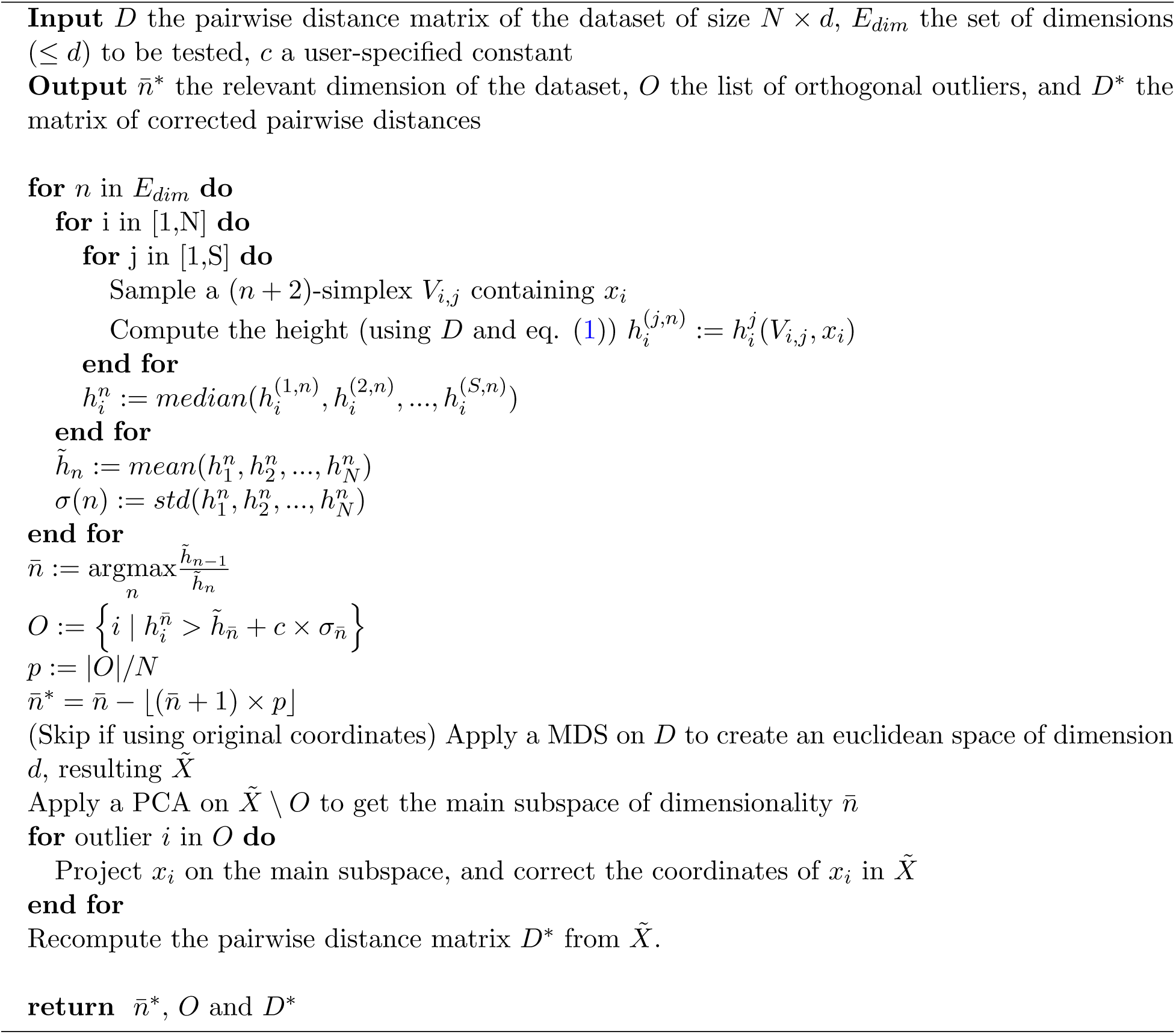

### 2.6 Datasets

The datasets of our study, described next, are available at this [OSF page](https://osf.io/x5796/).

#### Synthetic datasets

The “cross” dataset [5], which is a two-dimensional dataset representing a simple cross structure (Fig.2) was generated with *N* = 25 points, and *d*\* = 2. We introduced orthogonal outliers by randomly sampling three points and by adding a third coordinate of random amplitude to them. Other synthetic datasets were generated by sampling Gaussian-distributed coordinates in the main subspace, and adding some small noise in the whole space with variance between 0.0001 to 0.0003. A fraction *p* of the points was considered to define the orthogonal outliers, with coordinates modified by randomly increasing the coordinate(s) orthogonal to the plane; the amount increased is drawn from a uniform distribution between −30 and 30 or −100 to 100. These datasets were generated for a main subspace of dimension 2, 10 and 40, with *p* = 0.05 for dimensions *d*\* = 2 and 10, and *p* varying between 0.02 and 0.1 for *d*\* = 40. For all the synthetic datasets, the pairwise distance matrix was calculated using the Euclidean distance.

**Figure 2:**
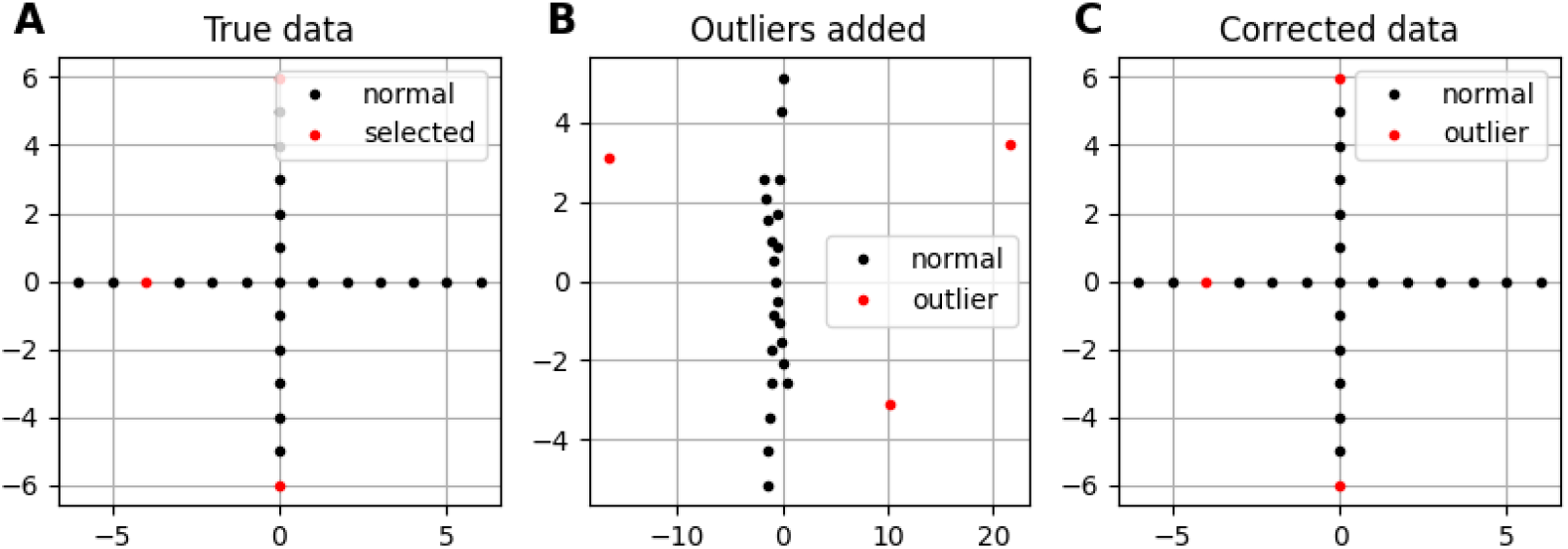
Application of DeCOr-MDS on a cross dataset. **A** Original cross dataset. the points selected to be orthogonal outliers are highlighted in red. **B** MDS embedding of the original data with an outlying component added to the selected points. **A** MDS embedding after preprocessing using DeCOr-MDS. Note that after correction, we recover the original cross structure.

#### Cell shape image dataset

The cell shape dataset contains mouse osteosarcoma 2D imaged cells [12], that were processed into a 100 × 2 vector of coordinates that define the cell shape contour, used as a test dataset in the Python package Geomstats [13] (for more details, see also [14] and the associated Github link). We more specifically considered the subset of “DUNN” cells (that denotes a specific lineage) from the control group (no treatment on the cells), which yields 207 cells in total. The pairwise distance matrix of all cell shapes was obtained from the same reference [13, 14] using the so-called Square Root Velocity metric that derives from the *L_2_* distance between velocities of the curves [15].

#### HMP dataset

The Human Microbiome Project (HMP) [16] dataset represents the microbiome measured across thousands of human subjects. The human microbiome corresponds to the set of microorganisms associated to the human body, including the gut flora, or the skin microbiota. The data used here corresponds to the HMP1 phase of clinical production. The hypervariable region v13 of ribosomal RNA was sequenced for each sample, which allowed to identify and count each specific microorganism, called phylotype. The processing and classification were performed by the HMP using MOTHUR, and made available as low quality counts (https://www.hmpdacc.org/hmp/HMMCP/) [16]. We downloaded this dataset, and subsequently, counts were filtered and normalized as previously described [10]. For our analysis, we also restricted our dataset to samples collected in nose and throat. Samples and phylogenies with less than 10 strictly positive counts were filtered out [10], resulting in an *n* × *p*-matrix where *n* = 270 samples and *p* = 425 phylotypes. Next, the data distribution was identified with an exponential distribution, by fitting its rate parameter. Normalization was then achieved by replacing the abundances (counts) with the corresponding quantiles. Lastly, the matrix of pairwise distances was obtained using the euclidean distance.

## 3 Results

### 3.1 Using n-simplices for orthogonal outlier detection and dimensionality reduction

We propose a robust method to reduce and infer the dimensionality of a dataset from its pairwise distance matrix, by detecting and correcting orthogonal outliers. The method, called *DeCOr-MDS*, can be divided into three sub-procedures detailed in sections 2.2-2.4), with the overall algorithm provided in section 2.5. The first procedure detects orthogonal outliers and estimates the subspace dimension using the statistics of simplices that are sampled from the data, using equations (2) and (3). The second procedure corrects for potential bias in estimated dimension when the fraction of outlier is large. The third procedure corrects the pairwise distance of the original data, by replacing the distance to orthogonal outliers by that to their estimated projection on the main subspace. The overall complexity, which we evaluate in details in Appendix (Supplementary file) is 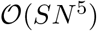, where *S* is the number of sampled simplices used in section 2.2, and *N* is the sample size of the data.

### 3.2 Performance on synthetic datasets

We first illustrate and evaluate the performance of the method on synthetic datasets, (for a detailed description of the datasets and their generation, see the Methods section 2.6). On a simple dataset of points forming a 2D cross embedded in 3D (Fig.2A), we observed that the MDS is sensitive to the presence of orthogonal outliers and distorts the cross when reducing the data in 2D (Fig.2B). In contrast, our procedure recovers the original geometry of the uncontaminated dataset, with the outliers being correctly projected (Fig.2C). The same results were obtained when sampling regular points from a 2D plane (Supplementary Fig.S1). We further tested higher dimensions, and illustrate in Fig.3A how the distribution of heights becomes concentrated around 0, when testing for the true dimension (*d*\* = 10), as suggested in the Methods section 2.2. As a result, our method allows to infer the main subspace dimension from Eq. (2), as shown in Fig.3B. In addition, the procedure accurately corrects the pairwise distances to orthogonal points with the distances to their projections on the main subspace, as shown in Fig.3C.

**Figure 3:**
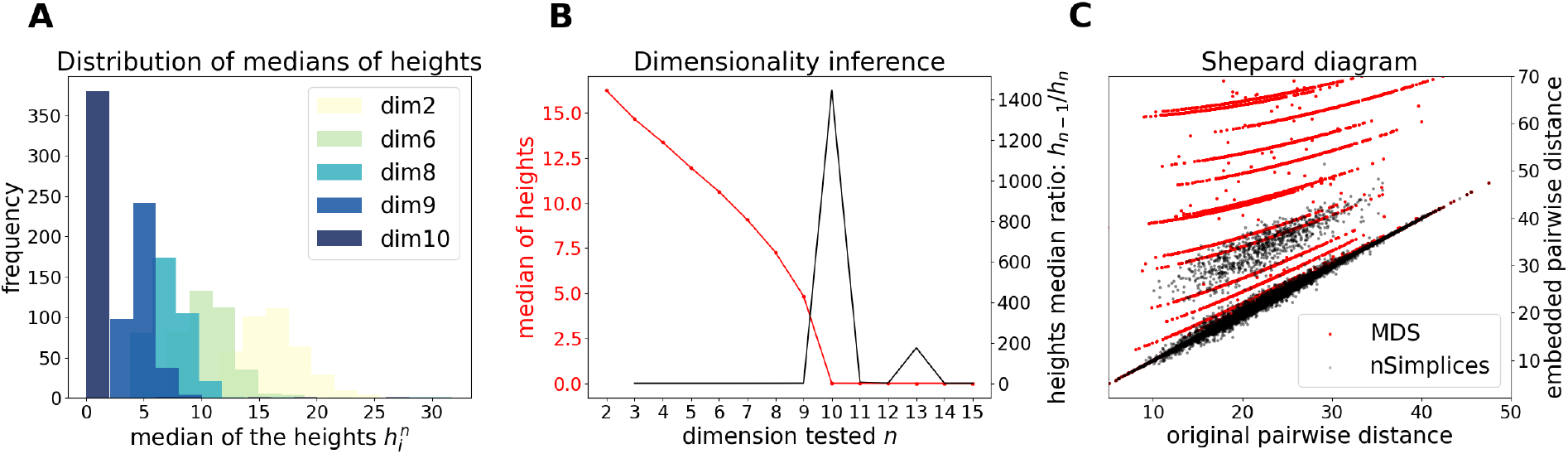
Application of DeCOr-MDS on a synthetic dataset with a main subspace of dimension 10. **A** Distribution of median heights per data point *x_i_* as a function of the tested dimension n. **B** Dimensionality inference based on the ratio of median heights (see also Eq. (1)), with the optimal ratio found for the true dimension 10. **C** Shepard diagram comparing the pairwise distances between regular points and outliers that are projected to the main subspace (true *δ_ij_*), with the same distances obtained after directly running MDS on the original pairwise distance matrix (red dots), or after correcting these distances using our procedure (black dots).

When the dimension of the subspace and fraction of outliers get significantly large, we finally illustrate the importance of the correction step (see Methods section 2.3), due to the sampling of simplices that contain several outliers. Upon using synthetic datasets with *d*\* = 40 and varying the fraction of outliers from 2% to 10%, we observe this bias appearing before correction, with *d*\* being overestimated by 2 or 3 dimensions (Fig.4). Using the debiased estimate n* from equation (4) successfully reduced the bias, with an error ≤ 1 for all the parameters tested.

**Figure 4:**
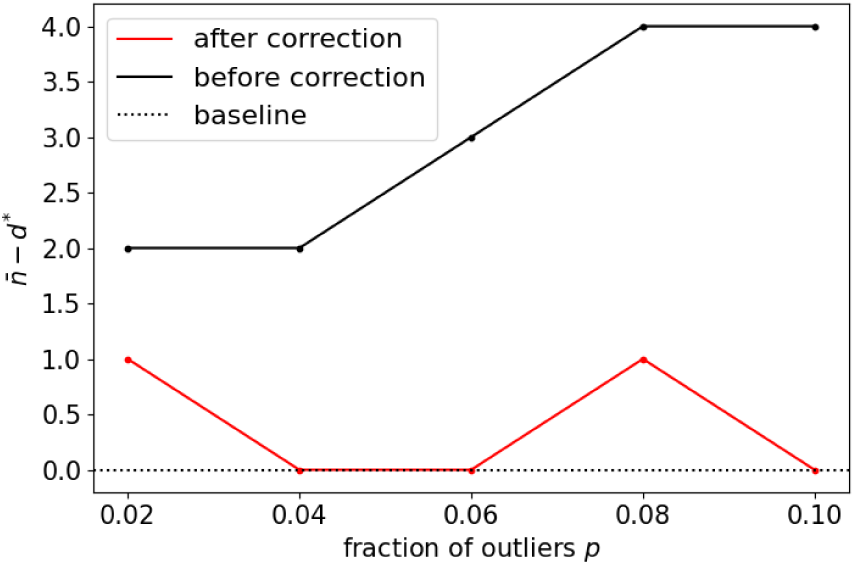
Application of DeCOr-MDS on a dataset with a main subspace of dimension 40: Dimension correction effect versus the fraction of outliers. The vertical axis represents the remaining bias between the inferred and actual dimensions, before and after bias correction. After correction, the differences between the estimated dimensions and the true dimension are always closer to 0 regardless of the fraction of outliers.

### 3.3 Application to cell shape image data

We further show how *DeCOr-MDS* can be broadly applied to biological data, ranging from images to high throughput sequencing. We first studied a dataset of single cell images, from osteosarcoma cells (see Fig.5A), which were processed to extract from their contour a 100 × 2 array of *xy* coordinates representing a discretization of a closed curve (see Dataset section 2.6). We obtained a pairwise distance matrix on this set of curves by using the so-called *Square Root Velocity* (SRV) metric, which defines a Euclidean distance on the space of velocities that derive from a regular parameterization of the curve [13, 15]. Using DeCOr-MDS, we found a main subspace of dimension 2 (Fig.5B), with 14 outliers detected among the 207 cells of this dataset. The comparison between the resulting embedding and that obtained from a simple MDS is shown in Supplementary Fig.S2, and reveals that outliers, when uncorrected, affect the embedding coordinates, while our correction mitigates it. By examining in more details the regular and inferred outlier cells (Fig.5C, with all cell shapes shown in Supplementary Fig.S3.), we found regular observations to approximately describe elliptic shapes, which is in agreement with the dimension found, since ellipses are defined by 2 parameters. One can also visually interpret the orthogonal outliers detected as being more irregular, with the presence of more spikes and small protusions. Interestingly, the procedure also identified as outliers some images containing errors, due to bad cropping or segmentation (with two cells shown instead of one), which should thus be removed of the dataset for downstream analysis.

**Figure 5:**
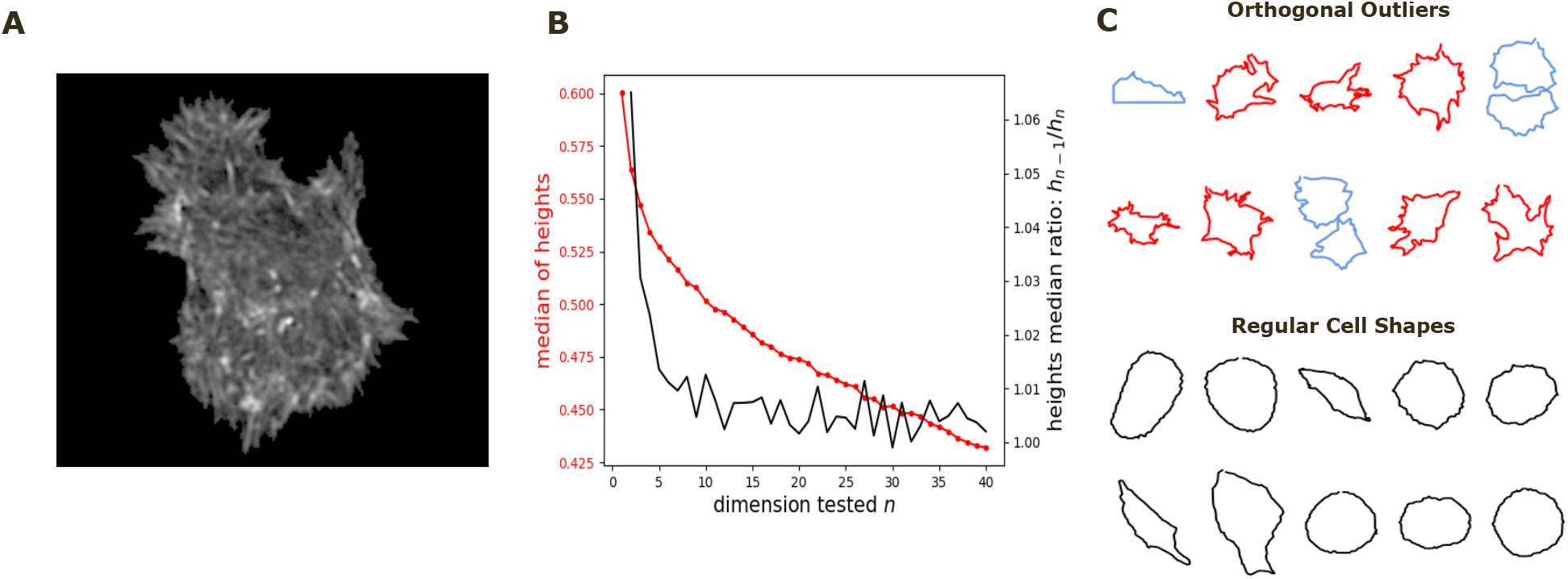
Application of DeCOr-MDS on a cell shapes dataset. **A**: An example of osteocarcoma cell image obtained from fluorescence microscopy. We process and extract the cell contour in our analysis. **B**: Dimensionality inference of the dataset obtained from 207 cell shapes using DeCOr-MDS. We estimate the dimension of the main subspace *n*\* = 2. **C**: Examples of cell shapes, including regular cells (in black), and orthogonal outliers detected. Among these outliers, we highlight cell shapes that are likely to be invalid due to segmentation errors (in blue), with the other outliers shown in red.

### 3.4 Application to HMP dataset

As another example of application to biological data, we next considered a dataset from the Human Microbiome Project (HMP). The Human Microbiome Project aims at describing and studying the microbial contribution to the human body. In particular, genes contributed by microbes in the gut are of primary importance in health and disease [16]. The resulting data is an array which typically contains the abundance of different elements of the microbiome (typically 10^2^ to 10^3^), denoted phylotypes, measured in different human subjects. To analyze such high dimensional datasets, dimensionality reduction methods including MDS (often denoted Principal Coordinates Analysis PCoA), are typically applied and used to visualize the data [17–19]. To assess our method incrementally, we restricted first the analysis to a representative specific site (nose), yielding a 136 × 425 array that was further normalized to generate Euclidean pairwise distance matrices (see Material and Methods section 2.6 for more details). Upon running *DeCOr-MDS*, we estimated the main dimension to be 3, with 6.62 % orthogonal outliers detected, as shown in Fig.6A. This is also supported by another study that the estimated dimension of HMP dataset is 2 or 3 [20]. We also computed the average distance between these orthogonal outliers and the barycenter of regular points in the reduced subspace, and obtained a decrease from 1.21 when using *MDS* to 0.91 when using *DeCOr-MDS*. This decrease suggests that orthogonal outliers get corrected and projected closer to the regular points, to improve the visualization of the data in the reduced subspace, like in our experiments with the synthetic datasets (Fig. 2 and Supplementary Fig. S2). In Fig.6B, we next aggregated data points from another site (throat) to study how the method performs in this case, yielding a 270 × 425 array that was further normalized to generate Euclidean pairwise distance matrices. As augmenting the dataset brings a separate cluster of data points, the dimension of the main dataset was then estimated to be 2, with 4.8 % orthogonal outliers detected, as shown in Fig.6B. The average distance between the projected outliers and the barycenter of projected regular points are approximately the same when using *MDS* (1.46) as when using *DeCOr-MDS* (1.45) for nose, and are also approximately the same when using *MDS* (1.75) to when using *DeCOr-MDS* (1.74) for throat. This decrease also suggests that orthogonal outliers get corrected and projected closer to the regular points.

**Figure 6:**
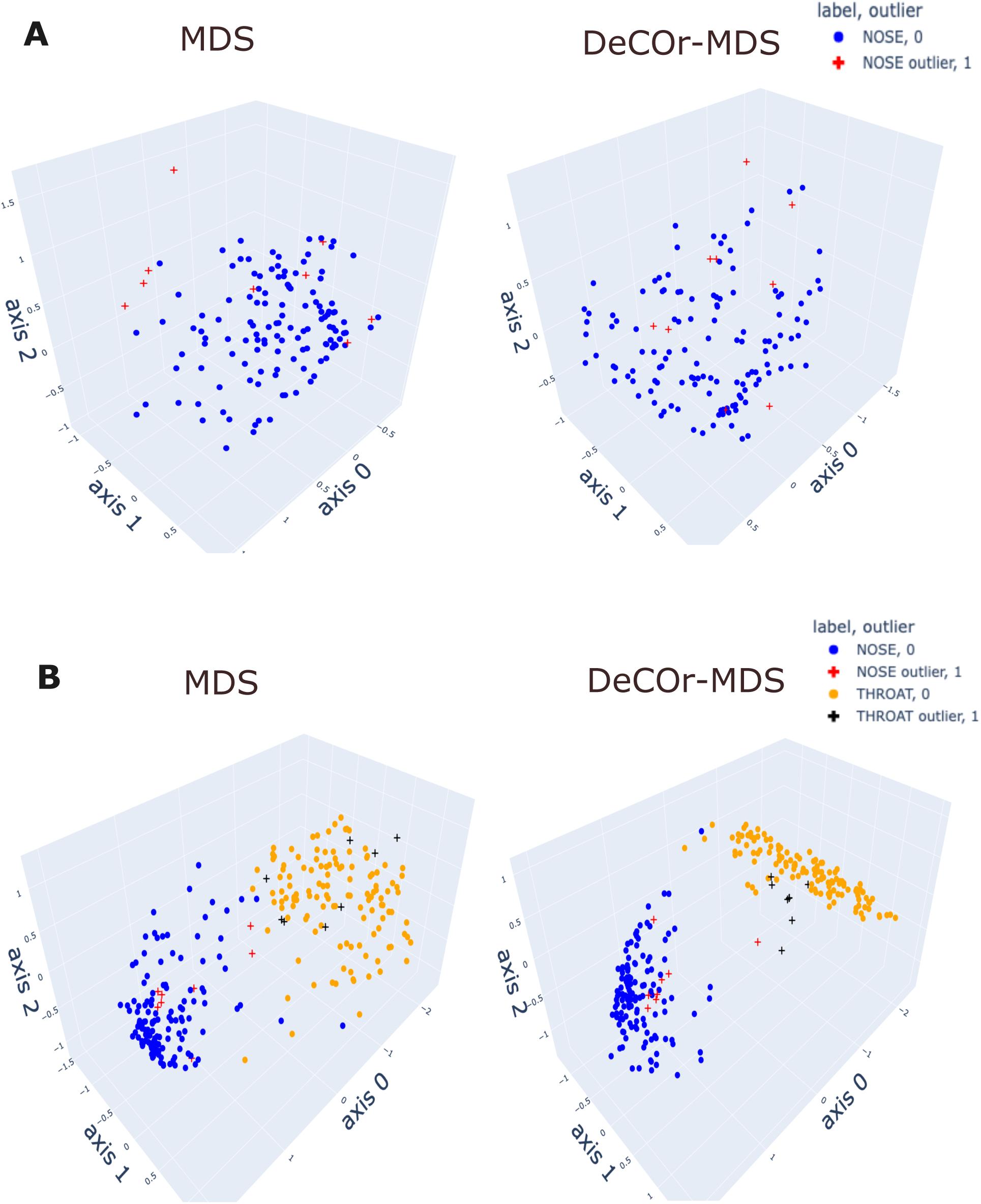
Application of DeCOr-MDS on HMP dataset. **A**: Structure restituted on 3 axes using *MDS* (left) and our procedure (right) using data from the nose site. The points marked with cross represent orthogonal outliers detected by *DeCOr-MDS*, which are also put closer to regular points after correction. **B** Same comparison as in (**A**) using data from nose and throat. The two clusters formed by nose and throat have a better seperation using DeCOr-MDS.

## 4 Discussion

We proposed *DeCOr-MDS*, a novel approach using geometric characteristics to detect dimension, and to correct orthogonal outliers in high dimensional space. That is, to the best of our knowledge, the first statistical tool that addresses the challenge of the presence of orthogonal outliers in high dimensional space. We validated the method using synthetic datasets and demonstrated its potential applications to analyze biological datasets, ranging from cell shape images to count arrays from microbiome data. The visualization and numerical comparison confirmed that *DeCOr-MDS* effectively detects dimensionalities in many instances, corrects orthogonal outliers, and demonstrates superior performance to classical dimension reduction methods.

The notion of simplices is used frequently with the aim of robustness, either to detect the coreness of data (data depth and multivariate median [21, 22]), or to detect outlying features (detection of extreme directions [23]). Simplices can also be used to build a flexible network of points for informative visualization [24]. Outlier detection and accommodation have been addressed by a wide array of methods, which can be broadly divided into three categories: (1) robust metrics [3–7], (2) robust estimation [8], or (3) exploiting the characteristics of outliers [2, 7]. Our method resorts to both (3) by using the geometry of data, and (1) by using the median as centrality estimator. Our method also aims at estimating dimension. A common approach to do so is the screeplot (or elbow) test in principal components analysis, where a notable drop in the proportion of variance (or distance) explained can be taken as a cutoff, and as the most relevant dimension. Highdimensional biological datasets challenge this strategy, because fine-scale structure confounds in practice downstream analyses. Because of this, authors often use an arbitrary large set of 10, or sometimes 20 or 50 components [25–30]. Power analyses based on simulations also provide a way to assess an adequate number of components [26]. In this work, we proposed an alternative approach, by exploiting the structure of the dataset to determine essential versus non-essential dimensions.

Limitations of DeCOr-MDS include the non-automated choice of the cutoff parameter c. This parameter sets the maximum tolerated number of standard deviations σ before a point is considered an outlier. A value for *c* = 3, which corresponds approximately to the 0.1% most extreme points in a normal distribution, may be selected, for instance. Although the method remained tractable for the datasets we analyzed, runtime complexity (analyzed in SI) is also significantly larger than for a standard MDS. Finally, dimension detection is still imperfect in datasets where the distribution of regular points (e.g. with distant clusters) may prevent the height criterion for outlier detection to be effective. There are various potential directions to improve the dimension detection in real datasets of high dimension. This may be achieved by studying the behaviour of the Cayley-Menger determinant, which is central in the procedure, in higher dimensions. One may also associate the height criterion with a distribution criterion [10], which would be sensitive to clusters or other notable structure, as was apparent in the HMP dataset. Another beneficial improvement would be to reduce computing time, for instance by implementing a parallelized version or using a call to a compiled program. Finally, one could optimize the cutoff parameter *c* automatically, either through a hyperparameter search, or by using a data-driven procedure, during the exploration phase of the algorithm.

## Acknowledgments

This research was supported by a NSERC Discovery grant (PG 22R3468) and a MITACS PIMS fellowship.

