## Supplemental Information for "Orthogonal outlier detection and dimension estimation for improved MDS embedding of biological datasets"

This Supplemental Information contains:

- Appendix section: Complexity of the procedure
- Supplementary Figures S1-3

### Appendix: Complexity of the procedure

The complexity of the algorithm can be evaluated as follows. Given one $n$-simplex, the volume computation is mainly a determinant computation, which has a complexity of $\mathcal{O}(n^3)$. The height computation of one point in one $n$-simplex does not add complexity. However, the height complexity, that relyes on computing the median of $S$ heights has a $\mathcal{O}(Sn^3)$ complexity. Thus, getting the heights of each point in the dataset amounts to $\mathcal{O}(SNn^3)$. The dimensionality and outliers detection corresponds to a maximum computation and applying a criterion to all the heights, which translates in a complexity of $\mathcal{O}(N)$. Finally, the complexity of the outlier correction reduces to the complexity of the PCA over the regular datapoints ($\mathcal{O}(Nd \times min(N, d) + d^3)$, the MDS over the pairwise distance matrix $D$ ($\mathcal{O}(N^3)$) and the final computation of the corrected distances matrix ($\mathcal{O}(dN^2)$). Thus, the complexity of the correction step is $\mathcal{O}(Nd \times min(N, d) + d^3 + N^3 + dN^2)$. Relying on a preliminary MDS for non-euclidean distances, the complexity of outlier correction becomes $\mathcal{O}(2N^3 + dN^2)$. Overall, if all dimensions are tested, the total complexity is $\mathcal{O}(SN^5 + N^2d + N^3 + d^3 + d^2N)$ (respectively, $\mathcal{O}(SN^5 + 2N^3 + dN^2)$ with preliminary MDS). Since $N > d$, the total complexity is $\mathcal{O}(SN^5)$.

#### Supplementary Figures

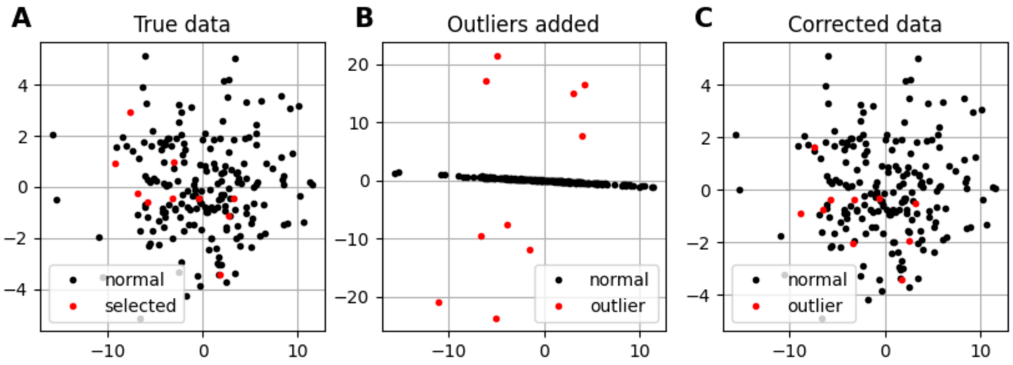


**Figure S1.** Application of DeCOr-MDS on a synthetic dataset with a main subspace of dimension 2. **A** MDS embedding of the original data; the points selected to be orthogonal outliers are highlighted in red. **B** MDS embedding of the data with an outlying component added to the selected points. **C** MDS embedding of the corrected data using DeCOr-MDS. Note that after correction, we recover the original structure.

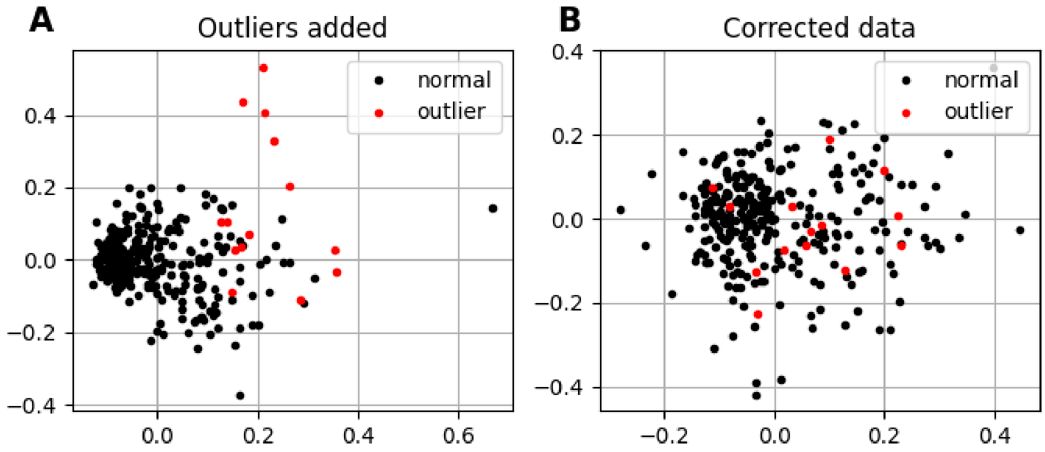


**Figure S2.** Application of DeCOr-MDS on a cell shape dataset with a main subspace of dimension 2. **A** MDS embedding of the data with an outlying component added to the selected points. **B** MDS embedding of the corrected data using DeCOr-MDS. We notice that the embedding in **A** is distorted, and outlier cells (red points) are corrected to be of the same magnitudes as normal cells.

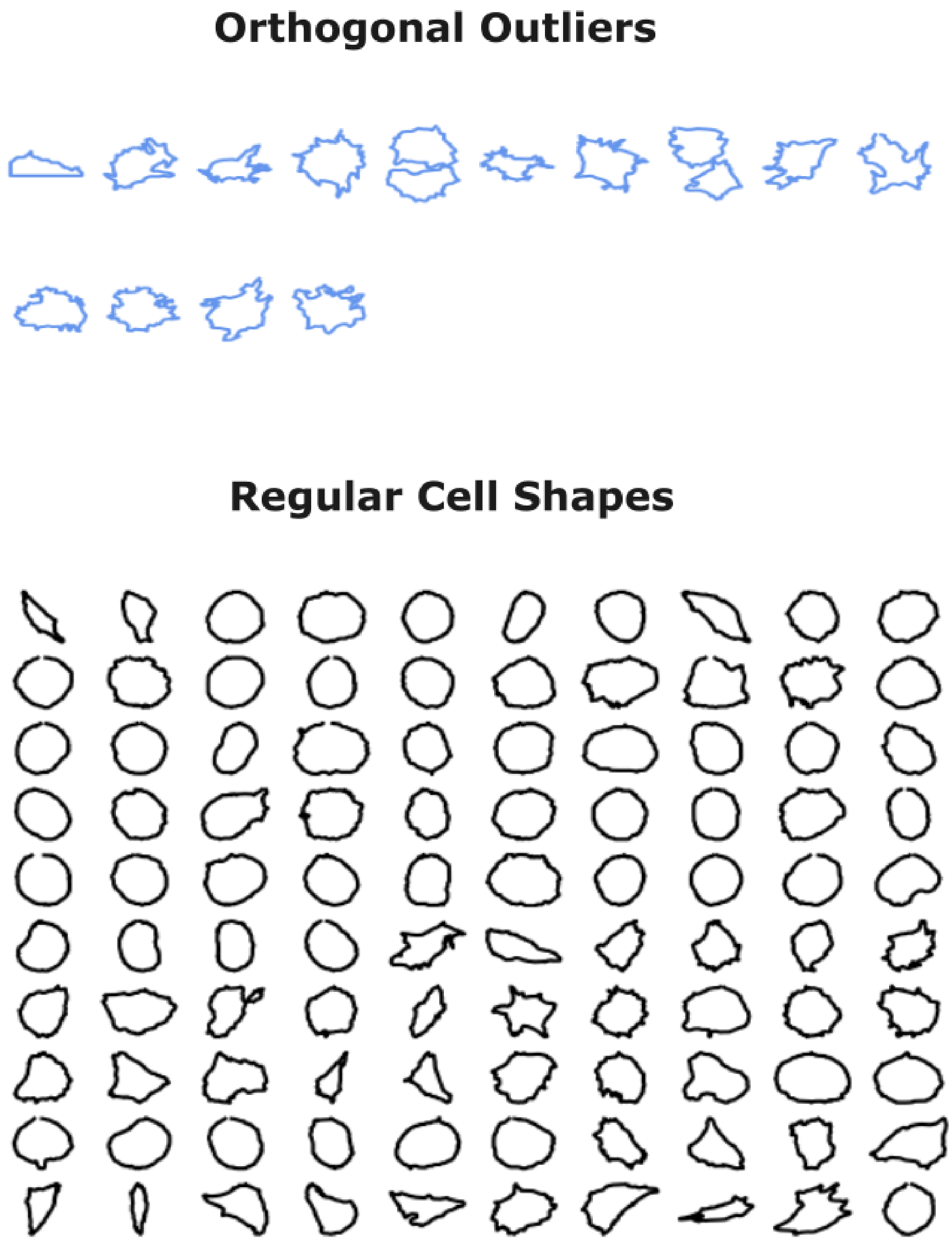


**Figure S3.** All 14 orthogonal outliers and the first 100 regular cell shapes identified by DeCOr-MDS.
